# ASSESSMENT OF CELL-TYPE-SPECIFIC EXCITATORY SYNAPTIC STRENGTH IN THE DORSOLATERAL STRIATUM OF GOAL-DIRECTED AND HABITUAL COCAINE-SEEKING BEHAVIOR

**DOI:** 10.1101/2025.10.07.680788

**Authors:** Kaliana M Veros, Maureen Timm, Peter J. West, Karen S Wilcox, Kristen A Keefe

## Abstract

With repeated exposure to addictive drugs, there is a shift from drug abuse to drug addiction that is mediated by the transition from goal-directed to habitual control. It is well known that the development of habitual control over behavior relies upon cell-type-specific synaptic changes in both D1 and D2 medium spiny neurons (MSNs) in dorsal striatum. Specifically, habitual behavior is mediated by increased synaptic strength in D2 MSNs in dorsolateral striatum (DLS), suggesting similar cell-type-specific synaptic changes may underlie the development of habitual cocaine-seeking behavior. However, cell-type-specific synaptic changes have not been evaluated in DLS in this context. Therefore, we trained rats to self-administer cocaine in a cocaine self-administration paradigm that allows for differentiation of goal-directed vs. habitual cocaine-seeking behavior. Moreover, we used a viral vector under a D2-specific promoter to fluorescently label D2 MSNs with eYFP in DLS. Evoked excitatory postsynaptic currents (EPSCs) were used to determine AMPA:NMDA receptor ratio and the rectification index. Surprisingly, we did not observe any significant differences in these measures in DLS of cocaine-seeking rats, regardless of whether cocaine seeking was under habitual control. Interestingly, preliminary observations revealed significant changes in the paired pulse ratio (PPR), suggesting that presynaptic mechanisms may be involved in the development of habitual control over cocaine seeking. Overall, however, these results suggest there are no changes in postsynaptic strength of D2 MSNs in the DLS of rats with an extended history of cocaine self-administration and regardless of whether the cocaine seeking is under goal-directed or habitual control.

**Significance Statement:** The study of drug abuse and drug addiction represents a critical area of research with significant public health implications. Importantly, the underlying neurobiology of the transition between drug abuse and drug addiction is not well understood and insights to this transition may aid in the development of novel treatment options. Behaviorally, the shift from goal-directed to habitual control is thought to underly this transition. Much is known about the neurobiology of goal-directed and habitual behavior, however the transition in the context of drug-seeking is not well defined. We observed no significant differences in measures of synaptic strength, suggesting such postsynaptic neuroplasticity in the dorsolateral striatum is not involved in this transition.

## Introduction

Drug addiction is a chronic relapsing disorder wherein drug-use behavior becomes inflexible and persistent even in the face of adverse consequences (Vanderschuren & Everitt, 2004; Belin et al., 2013). Initially, drugs are obtained and consumed to intentionally experience a reinforcing outcome, and drug-seeking behaviors are thereby outcome-motivated and flexible (Ersche et al., 2016; Vanderschuren & Everitt, 2004). As drug use continues, the behaviors involved in drug consumption become automatic in response to drug-associated stimuli (Ersche et al., 2016; Everett & Robbins, 2005). As such, drug-related behavior is considered to undergo a shift from goal-directed to habitual performance (Ersche et al., 2016; Pierce & Vanderschuren, 2010). Therefore, understanding behavioral and neurobiological processes involved in the progression of drug-related behavior from goal directed to habitual is critical.

Goal-directed and habitual instrumental behaviors are known to be mediated by distinct subregions within dorsal striatum--the dorsomedial and dorsolateral striatum (DMS and DLS), respectively (Packard & Knowlton, 2002; Yin et al., 2004; 2006; 2009; Dolan & Dayan, 2013; Lipton et al., 2019). Importantly, the development of habitual control over appetitive instrumental behavior is associated with changes in glutamatergic synapses of DLS (Lipton et al., 2019; Malvaez, 2020). Early studies demonstrated increases in excitatory synaptic strength with extended training, suggesting that potentiated glutamate signaling in DLS underlies habitual actions (Yin et al., 2009). Further, as medium spiny neurons (MSNs) in dorsal striatum are segregated into dopamine D1 receptor or D2 receptor-expressing MSNs (Calabresi et al., 2014), other studies examined cell-type-specific changes in DLS (Lovinger et al., 2010; Shan et al., 2014; O’Hare et al., 2016; Lipton et al., 2019). Specifically, D2 MSNs in DLS reportedly exhibit synaptic strengthening in comparison to D1 MSNs after extended training, suggesting synaptic potentiation in D2 MSNs may underlie habitual control over behavior (Yin et al., 2009; O’Hare et al., 2016). While these studies indicate cell-type-specific changes in DLS in relation to habitual control over appetitive instrumental behavior, little is known about cell-type-specific changes in MSNs of DLS associated with habitual drug-seeking behavior.

In cocaine self-administration paradigms, inactivation of DLS reverts habitual cocaine-seeking behavior to goal-directed, suggesting drug-related behaviors are similarly mediated by dorsal striatum (Zapata et al., 2010; Jones et al., 2022). Despite these observations, cell-type-specific mechanistic changes within DLS underlying habitual drug-related behaviors remain poorly defined. Mechanistic changes have been well studied in the nucleus accumbens (NAc) in the context of cocaine self-administration (Kalivas, 2009). Most notably, persistent cocaine craving is associated with insertion of inwardly rectifying Ca^2+^-permeable AMPA (CP-AMPA) receptors in the NAc, enhancing MSN excitability and strengthening synapses (Wolf, 2016). Moreover, D1 MSNs exhibit CP-AMPA receptor insertion following 10 days of cocaine self-administration (Terrier et al., 2016; Zinsmaier et al., 2021). However, given the importance of DLS in habitual control over behavior, cell-type-specific changes in the context of habitual cocaine-seeking represent a critical area of study in understanding synaptic changes underlying transition from drug abuse to drug addiction.

Therefore, the objective of the present study was to determine if there are changes in glutamatergic synaptic transmission in DLS of rats characterized as goal-directed or habitual in their cocaine-seeking behavior. Rats were assessed for sensitivity to outcome devaluation following training on a chained cocaine self-administration paradigm to determine goal-directed or habitual control over cocaine seeking (Zapata et al., 2010; Giangrasso et al., 2023). We then assessed synaptic strength in DLS via patch-clamp recordings of evoked excitatory postsynaptic currents (EPSCs) in cells expressing eYFP under a D2 dopamine receptor-specific promotor (i.e., D2 MSNs), as well as in unlabeled cells (i.e. D1 *or* D2 MSNs). In all cell types, we found that excitatory synaptic strength, assessed through AMPA:NMDA receptor ratio and rectification index (RI), was unchanged in rats characterized as goal-directed or habitual relative to yoked-saline controls. Interestingly, we did observe a statistically significant decrease in the paired pulse ratio (PPR) of EPSCs in eYFP+ cells from habitual rats as compared to rats characterized as being goal-directed in their cocaine-seeking, as well as a significant increase in PPR of EPSCs in eYFP-MSNs relative to saline controls, indicating presynaptic release may be differentially altered onto MSN subpopulations. Overall, the present results suggest no change in excitatory synaptic input onto D2 or putative D1 MSNs in DLS of rats with a history of cocaine self-administration, regardless of whether the cocaine seeking is under goal-directed or habitual control. Future studies thoroughly interrogating presynaptic release properties onto DLS MSNs are warranted, however, in light of the present preliminary findings of altered PPRs in rats with habitual control over their cocaine-seeking behavior.

## Materials and Methods

### Animals

Male Long Evans rats (300–350 g) surgically prepared with right jugular vein catheters were obtained from Charles River Laboratories (Wilmington, MA, United States). Upon arrival, catheter patency was verified and maintained by daily infusions of heparin-dextrose catheter-locking solution (500 IU/50% dextrose; SAI Infusion, IL, United States) and daily infusions of prophylactic Baytril (10mg/kg, i.v.; Norbrook, Newry, United Kingdom). All rats underwent stereotaxic surgery 6–8 days after arrival and were allowed to recover for 5–7 days before the start of behavioral training. Rats were single-housed in standard housing conditions following surgery to prevent the rats from damaging the indwelling catheter and were randomly assigned to receive either cocaine training or to serve as yoked-saline controls. Three days before the start of cocaine self-administration training, rats were food-restricted to 25 g of standard chow per day, fed following training, with *ad libitum* access to water. Rats were maintained on this feeding schedule throughout experimental training. Experimental training was performed between ZT3 and ZT7. Animal care, surgeries, and experimental procedures followed the Guide for the Care and Use of Laboratory Animals (8^th^ Edition) and were performed in accordance with the [Author University] animal care committee’s regulations.

### Stereotaxic surgery

Rats were anesthetized with isoflurane (2.0–2.5% induction; 1.5–2.0% maintenance) and placed in a stereotaxic apparatus (Stoelting, IL, United States). A small hole was drilled in the skull and a 28-gauge infusion cannula (P1 Technologies, VA, United States) was inserted into the DLS (in mm relative to bregma: AP +0.7, ML −3.6, DV −5.0; Yin et al., 2004; Giangrasso et al., 2023). In order to identify dopamine D2 receptor-expressing MSNs for patch clamp studies, a total of 1μL of AAV-8-D2SP-eYFP-WPRE (Stanford Gene Vector and Virus Core, GVVC ID: GVVC-AAV-114; Zalocusky et al., 2016) was infused into DLS over the course of 10 min, after which the cannula was left in place for an additional 5 min before removal to ensure maximal diffusion. Rats were given 10 mg/kg Baytril i.v. and 0.1 mL of heparin-dextrose catheter-locking solution (500 IU/50% dextrose; SAI Infusion, IL, United States) on the day of surgery. Rats also were given 5 mg/kg carprofen s.c., for prophylactic pain management on the day of and for three days after surgery. Throughout the recovery period and cocaine self-administration training and testing, rats were given daily infusions of i.v. Baytril and heparin-dextrose catheter-locking solution following training. Rats were allowed to recover from stereotaxic surgery for 3–5 days before food restriction began, and 5–7 days before the start of self-administration training.

### Cocaine self-administration paradigm

The cocaine self-administration paradigm, adapted from Zapata et al. (2010) and Giangrasso et al. (2023), was performed during the light cycle and conducted in eight standard operant chambers that were enclosed in sound and light-attenuating cabinets (Coulbourn Instruments, PA, United States). One wall of the chamber had two retractable levers on the right and left sides, and the opposite wall was equipped with a 3-W, 24-V house light. Graphic State 4.0 software (Coulbourn Instruments, PA, United States) was used to control chamber equipment and experimental protocols, as well as to record the number of lever presses and session time. Rats were trained to press one lever (designated as the drug “taking” lever) for intravenous cocaine•HCl infusion (0.33 mg/50 μL infusion; calculated as the salt; NIDA Drug Supply Program, NC, USA) under a fixed ratio 1 (FR1) schedule. Each taking-lever response was accompanied by retraction of the taking lever, illumination of a cue light, and extinction of the house light for a 30 second time-out (TO) period. Rats were trained daily on the taking lever for 2 h or 40 infusions per session, whichever came first. A criterion of 2 consecutive sessions with >10 cocaine infusions per session was required before rats progressed to a chained seeking-taking schedule. In this phase of training, an additional, different lever, designated as the “seeking” lever, was introduced. The first press of the seeking lever, under a 2-second random interval (RI) schedule, resulted in retraction of the seeking lever and presentation of the taking lever. Responding on the taking lever was kept on the FR1 schedule of cocaine administration, followed by the 30 second TO period. Following the TO period, the seeking lever was reinserted, and the chained cycle began again. This seeking-taking chained schedule is denoted as: RI(2s)/FR1:TO(30s) or “Chain #1.” Rats underwent daily training on this schedule (3 h or 12 infusions, whichever came first) until they reached criterion of two consecutive sessions with 12 infusions/session. Rats then progressed through increasing chained schedules: Chain #2 RI(20s)/FR1:TO(120 s) and Chain #3 RI(60s)/ FR1:TO(300 s) using the same training criteria. They then advanced to Chain #4 RI(120 s)/FR1:TO(600 s), which continued for six sessions (12 infusions/session), at which point all rats had achieved stable responding (<20% variability in drug infusions). A subset of rats underwent training on Chain #4 for six additional days to promote habitual control over behavior. Yoked-saline controls received response-independent saline infusions identical to that of their paired experimental rat and lever-pressing had no scheduled consequences.

Upon completion of Chain #4, rats progressed to 13 days of outcome devaluation wherein only the taking lever was available, and no cocaine was delivered upon lever-pressing, thereby devaluing the drug-taking lever via outcome omission. After the last day of outcome devaluation, seeking behavior was assessed under these “devalued” conditions in a 5-minute test session. In this seeking test, only the seeking-lever was available and responding did not result in any consequences. Following the “devalued” seeking test, the taking lever was then re-valued over two daily sessions identical to the initial FR1 taking lever training sessions (2h or 40 infusions, whichever came first). The following day, another 5-minute seeking test was conducted under these “valued” conditions. Comparing the number of cocaine-seeking lever presses made under the “devalued” vs. the “valued” test conditions gives insight into the sensitivity of drug-seeking to outcome devaluation, thereby enabling the classification of cocaine-seeking behavior as either under goal-directed or habitual control (Giangrasso et al., 2023; Zapata et al., 2010). A “cocaine-seeking score” was determined for each rat as the number of seeking responses made under the “devalued” test condition expressed as a percentage of seeking responses made under the “valued” test condition. Rats with cocaine-seeking scores ≤70% were classified as having “goal-directed” cocaine-seeking behavior, scores between 71% – 79% as “intermediate,” and scores ≥80% were classified as “habitual,” similar to that previously reported by others (Zapata et al., 2010; Giangrasso et al., 2023). All rats were sacrificed within 1–4 days following completion of the final seeking test for electrophysiology recordings. We limited this time course to 1–4 days after the testing as work on incubation of cocaine craving showed no significant changes in drug seeking over days 1–4 (Grimm et al., 2001) and subsequent studies of incubation variably use 1–4 days as “early” withdrawal time points (Grimm et al., 2001; Lu et al., 2004). The 1–4-day period should, therefore, have mitigated potential confounding effects of withdrawal and incubation of craving on the dependent measures.

### Brain slice preparation

From the point of brain slice preparation onward through final data analysis, the experimenter was blinded to the treatment group and behavioral classification of the rats. Within 4 days after the final seeking test, rats were anesthetized with sodium pentobarbital (50 mg/kg, i.p.) and immediately decapitated. The brain was then divided into two hemispheres longitudinally. Coronal brain slices (350 μm) containing the DLS were collected in an oxygenated ice-cold NMDG-HEPES cutting solution (in mM: 92 NMDG, 2.5 KCl, 1.2 NaH_2_PO4, 30 NaHCO_3_, 20 HEPES, 25 glucose, 2 thiourea, 5 Na-ascorbate, 3 Na-pyruvate, 10 MgSO_4_, and 0.5 CaCl_2_). Slices were then transferred to a pre-warmed (32–24°C) holding chamber containing NMDG-HEPES for 30 minutes as a protective recovery period. Stepwise, a Na+ spike-in procedure was carried out according to an optimal age-dependent schedule (Ting et al., 2014). After 30 minutes, slices were transferred to a holding chamber containing room-temperature oxygenated HEPES-aCSF holding solution (in mM: 92 NaCl, 2.5 KCl, 1.2 NaH_2_PO_4_, 30 NaHCO_3_, 20 HEPES, 25 glucose, 2 thiourea, 5 Na-ascorbate, 3 Na-pyruvate, 2 MgSO_4_, and 2 CaCl_2_). All solutions were bubbled with 95% O_2_/5% CO_2_ throughout the experiment, corrected to pH 7.30– 7.35, and adjusted for an osmolarity between 295–300 mOsm.

### Electrophysiology

Whole-cell patch clamp recordings of MSNs were obtained by recording in the voltage-clamp configuration through use of a Multiclamp 700B amplifier, a Digidata 1440A data acquisition board, and pClamp10 software (Molecular Devices, CA, USA). Slices were visualized with a 40x water-immersion objective (NA 0.8; Carl Zeiss, Thornwood, NY) using infrared differential interference contrast (IR-DIC) microscopy on an upright Axioskop2 microscope (Carl Zeiss, Thornwood, NY) prior to specific MSN identification. Cells displaying eYFP fluorescence (excitation 513 nm, emission 527 nm) deep in the tissue were targeted as eYFP+ cells for electrophysiology with patch pipettes containing a CsMeSO_3_ intracellular solution (in mM: 120 CsMeSO_3_, 15 CsCl, 8 NaCl, 10 HEPES, 0.2 EGTA, 10 TEA, 5 QX-314, 0.1 Spermine, 4 MgATP, and 0.3 NaGTP; pH 7.28–7.33; osmolarity 290–295 mOsm). Cells not displaying eYFP fluorescence were targeted as eYFP-cells for electrophysiology. Resting membrane potential was determined in the *I=0* mode, and only cells with hyperpolarized resting membrane potentials (< −65mV) were included in analyses. Membrane and access resistance were monitored, and only stable recordings were included in the study.

A bipolar nichrome/formvar stimulating electrode placed 100–200 μm from the patch pipette in DLS was used to deliver stimulation to evoke EPSCs. A maximum of two MSNs (one eYFP+, one eYFP-) were patched per slice. Signals were acquired at 10 kHz and filtered at 2 kHz. Brain slices were recorded in oxygenated aCSF at 30°C (in mM: 126 NaCl, 2.5 KCl, 1 NaH_2_PO_4_, 26 NaHCO_3_, 10.5 glucose, 1.3 MgSO_4_, 2 CaCl_2_) containing picrotoxin (50μM). EPSCs at holding potentials ranging from −70mV through +40mV were evoked using a stimulation intensity that elicited response amplitudes from 100–300pA at −70mV (0.4mA – 1mA). Evoked EPSC current-voltage relationships were determined by normalizing peak amplitude at each holding potential to the first evoked response at −70mV. RI was determined as the ratio of peak EPSC amplitudes recorded at −70mV to those recorded at +40mV and used as a measure of the insertion of CP-AMPA receptors into the neurons. AMPA:NMDA receptor ratio was determined as the ratio of peak EPSC amplitudes recorded at −70mV to peak amplitude 50–60ms after stimulation at +40mV (Shan et al., 2014). PPRs were determined as the ratio of peak amplitude of the second pulse to that of the first at −70mV with an inter-stimulus-interval (ISI) of 50ms. All electrophysiological analyses were performed on baseline-corrected traces.

### RNAScope/immunohistochemistry

For verification of viral eYFP expression in D2 MSNs, a combination of RNAScope™ and immunohistochemistry (IHC) was used. Rats were anesthetized with sodium pentobarbital (50 mg/kg, i.p.), immediately decapitated, and the brain was flash frozen in 2-methylbutane (MilliporeSigma, MA, United States) and stored at −80°C until use. Brains were cryosectioned at 14 μm and thaw mounted on Superfrost® Plus slides (VWR International). Coronal sections containing the dorsal striatum were processed using RNAScope™ (Advanced Cell Diagnostics a Bio-Techne Brand, CA, United States) following the manufacturer’s protocol with probes against preproenkephalin (Penk1; RNAScope™ Probe-Rn-Penk, Cat No. 417431) as a marker for D2 MSNs or preprotachykinin (Tac1; RNAScope™ Probe-Rn-Tac1, Cat No. 450661) as a marker for D1 MSNs. Immediately following the RNAScope™ protocol, slides were incubated in blocking solution (0.1% Triton X-100, 10% normal goat serum in PBS) for 2 hours. Slides were then incubated overnight in primary anti-GFP antibody (1:500 dilution, Cat No. AB6556; Abcam; RRID:AB_305564) in blocking solution (0.1% Triton X-100, 1% normal goat serum in PBS). The primary antibody was visualized using an Alexa Fluor 488-conjugated secondary antibody (1:1000 dilution, 0.1% Triton X-100, 1% normal goat serum in PBS, Cat No. A11008; Invitrogen; RRID:AB_143165). Slides were coverslipped using ProLong™ Diamond with DAPI (Cat No. P36971; Invitrogen). Areas of the DLS expressing GFP were first identified at 20x magnification. A confocal z-stack (10 μm; 1-μm z-steps) of the DLS containing GFP-expressing cells was then acquired at 63x magnification using a Zeiss LSM700 laser scanning confocal microscope (Carl Zeiss, Oberkochen, Germany).

### Statistical analysis

All statistical analyses were performed in GraphPad Prism (GraphPad Software Inc., CA, United States). For all analyses, alpha was set to 0.05 and power analyses were conducted with a desired power level of 0.8. D-Agostino & Pearson normality tests were performed on all data sets to determine if the data were normally distributed. If the data were normally distributed (*p* > 0.05), parametric tests were used, including: one-way ANOVA and unpaired t-tests. If the data were not normally distributed (*p* < 0.05), nonparametric versions of the aforementioned tests were used, including: Kruskall Wallis test, Mann–Whitney U test, and Spearman correlation. As noted above, all electrophysiology recordings and data analyses were conducted with the experimenter blinded to the treatment and behavioral classification of the subject.

## Results

### Cocaine-seeking behavior

We used a chained cocaine self-administration paradigm, adapted from Zapata et al. (2010) and Giangrasso et al. (2023) to train male rats (*n* = 32 rats total) in cocaine-seeking and -taking behaviors. Through devaluation of drug-taking responses via outcome omission, the number of seeking-lever responses made under devalued and valued conditions were used to calculate the cocaine-seeking score and to classify cocaine-seeking behavior as either goal-directed (cocaine-seeking score ≤ 70%; *n* = 17), intermediate (cocaine-seeking score 71–79%; *n* = 3), or habitual (cocaine-seeking score ≥ 80%; *n* = 12) (**Figure 1**). All rats readily met criteria for cocaine-taking responses in initial training with a total average time to meet criteria on FR1 of 3.75 ± 0.42 days. The average number of days to meet criteria on the FR1 was similar between groups classified as goal-directed (4 ± 0.65 days; *n* = 17), intermediate (3.33 ± 1.33 days; *n* = 3), or habitual (3.5 ± 0.57 days; *n* = 12) (one-way *ANOVA*: F_(2, 29)_ = 0.199, *p* = 0.82^a^). All rats reduced responding during devaluation training, with no significant differences observed on the final day of devaluation training between goal-directed (13.1± 2.56 lever presses; *n =* 17), intermediate (29.3 ± 9.67 lever presses; *n =* 3), or habitual (49.5 ± 28.1 lever presses; *n =* 12) classifications (*Kruskal-Wallis statistic* = 4.625; *p =* 0.099^b^). Further, all rats readily reinstated cocaine-taking behavior in revaluation training sessions (22.8 ± 1.1 lever presses) to rates similar to those observed in initial FR1 training (20.8 ± 1.37 lever presses; *Mann Whitney U =* 373; *p =* 0.061^c^). A subset (*n* = 10) of the animals referred to above were run on Chain #4 for an additional six days (i.e., Chain #4 for a total of 12 days) to promote habitual responding. Within this cohort, few remained goal-directed (*n* = 2) or intermediate (*n* = 2), whereas the majority were classified as habitual (*n* = 6) in their cocaine-seeking behavior. Following electrophysiology recordings and exclusions, the intermediate group was omitted from further analysis for lack of sufficient group size. Therefore, analysis of all electrophysiological data was performed on cells recorded in brain slices obtained from yoked-saline control rats (*n* = 7 rats) and cocaine self-administering rats classified as goal-directed (*n* = 11) and habitual (*n* = 10) in their cocaine-seeking behavior.

**Figure 1:**
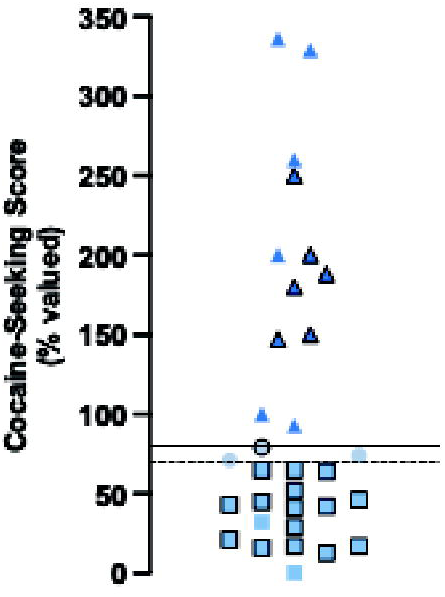
Cocaine-seeking scores of rats subjected to chained cocaine-seeking and - taking self-administration training. Distribution of cocaine-seeking scores *(n* = 32 rats; goal-directed: *n* = 17; intermediate: *n* = 3; habitual: *n* = 12). Cocaine-seeking scores (seeking responses under “devalued” conditions expressed as a percentage of responses under “valued” conditions) are classified as goal­directed (≤70%, squares), intermediate (71% - 79%, circles), and habitual (≥80%, triangles). Data points without borders represent rats trained an additional six days on Chain #4 *(n* = 10). Dashed line represents cocaine-seeking score of 70%, data points below this line are considered goal-directed. Solid line represents cocaine-seeking score of 80%, data points above this line are considered habitual.

### Baseline electrophysiological characteristics of eYFP+ and eYFP-cells

To assess synaptic strength, we recorded stimulation-evoked excitatory postsynaptic currents from both eYFP+ and eYFP-neurons in the DLS of acute brain slices obtained from yoked-saline controls and cocaine-experienced rats. We confirmed that eYFP expression was selective for D2 MSNs, rather than D1 MSNs, through RNAScope™/IHC analysis of Penk1 and Tac1 mRNAs and eYFP protein expression. Importantly, Penk1 is a known marker of MSNs expressing D2 receptors, whereas Tac1 is known to be expressed in D1 MSNs throughout the striatum (Gerfen et al., 1990; Pollack & Wooten, 1991). RNAScope™/IHC analysis revealed colocalization of Penk1/eYFP and lack of colocalization of Tac1/eYFP (**Figure 2**), thereby suggesting that the viral eYFP expression is restricted to D2 MSNs.

**Figure 2:**
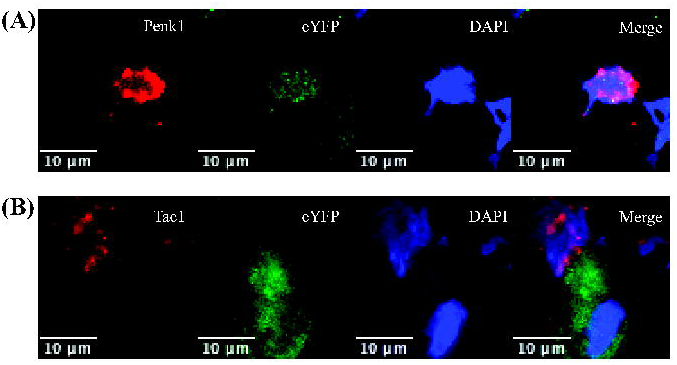
eYFP expression in identified D2 MSNs in the DLS. **(A)** A representative cell double-labeled for preproenkephalin (Penk1; red; D2 MSN) and eYFP (green) with DAPI counterstain (blue) in the DLS. **(B)** A representative image showing eYFP expression (green) is *not* colocalized with preprotachykinin (Tac1; red; D1 MSN) expressing cells.

In electrophysiological recordings, intrinsic properties were measured to ensure recordings were obtained from MSNs. MSNs were differentiated from other cell types (e.g., cholinergic interneurons) by a hyperpolarized resting membrane potential (−80 to −90mV) and low input resistance (<100MΩ; Kreitzer & Malenka, 2008; Cepeda et al., 2008). Importantly, baseline characteristics (**Table 1**) were not different between voltage-clamped eYFP+ (*n* = 28 cells) and eYFP-(*n* = 21 cells) cells, including: resting membrane potential (*unpaired t-test; p =* 0.93^d^); input resistance (*unpaired t-test; p =* 0.35^e^) and evoked EPSC stimulation strength *(unpaired t-test; p =* 0.59^f^). Together, these measures suggest that eYFP+ cells were D2 MSNs, that eYFP-cells were D1 MSNs or non-transfected D2 MSNs, and that eYFP expression does not influence electrophysiological characteristics of the recorded MSNs.

### Analyses of electrophysiological recordings reveal no changes in excitatory synaptic strength in MSNs expressing eYFP

Analyses of EPSCs are common in the assessment of synaptic strength. Evoked EPSCs at a range of holding potentials provides insight to the relative distribution of AMPA and NMDA receptors, as well as the general current-voltage relationship of postsynaptic neurons. Therefore, we recorded evoked EPSCs in voltage-clamped MSNs in the DLS of acute brain slices obtained from yoked-saline controls and rats classified as goal-directed or habitual in their cocaine-seeking behavior. Evoked EPSCs were recorded at a range of voltages (−70mV – +40mV) to assess changes in various synaptic parameters. In D2 MSNs there were no apparent differences in the current-voltage relationship of evoked EPSCs between saline and cocaine-experienced rats when comparing normalized EPSCs at holding potentials between −70 and +40 mV (**Figure 3B**). Likewise, no differences were apparent when recordings obtained from cocaine-experienced rats were broken down based on their behavioral classification as either goal directed or habitual in their cocaine seeking. (**Figure 3A,B**).

**Figure 3:**
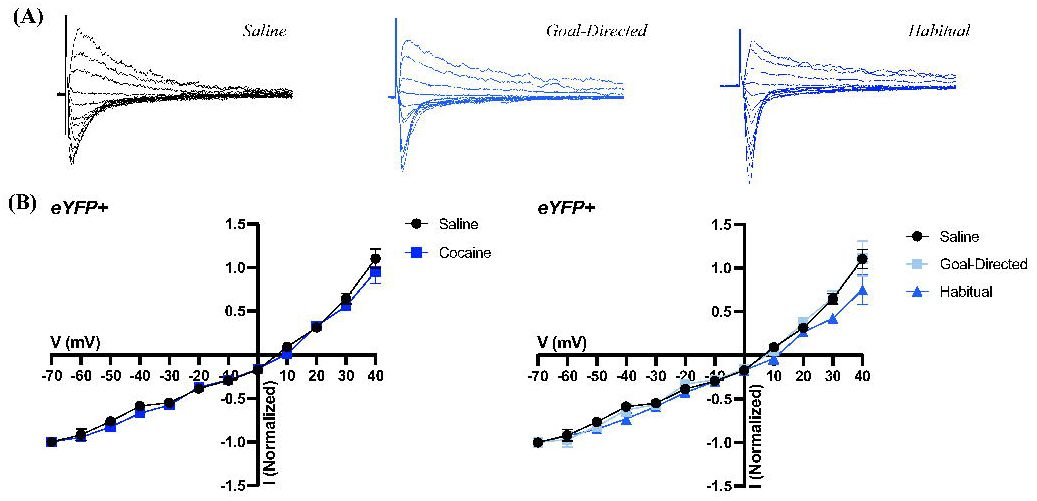
EPSCs in eYFP+ cells of the DLS obtained from acute brain slices of cocaine­ experienced and yoked-saline control rats. **(A)** Representative normalized traces of evoked EPSCs in brain slices from control (left) and cocaine-experienced rats classified as goal-directed (middle) or habitual (right) in their cocaine­ seeking behavior. **(B)** IN relation of evoked EPSC responses between −70mV and +40mV in the DLS of eYFP+ cells between saline *(n* = 8 cells) and cocaine-experienced rats (left; *n* = 19 cells) classified as goal-directed *(n* = 11 cells) or habitual *(n* = 8 cells) in their drug-seeking behavior (right).

The insertion of inwardly rectifying CP-AMPA receptors has been demonstrated in the NAc following extended cocaine self-administration and withdrawal and is associated with incubation of cocaine craving (Wolf, 2016). Therefore, we hypothesized that insertion of CP-AMPA receptors in the DLS following cocaine exposure may similarly contribute to the development of habitual control over cocaine-seeking behavior. To assess whether there might be differences in the insertion of CP-AMPA receptors in the DLS of rats whose cocaine-seeking behavior was habitual, we examined the RI from evoked EPSCs. As CP-AMPA receptors are inwardly rectifying, an increase in the RI is interpreted as indicative of insertion of CP-AMPA receptors into the cell membrane. However, analyses of the RI did not reveal any differences in eYFP+ cells of the DLS from yoked-saline controls (0.99 ± 0.13; *n* = 8 cells) vs rats with a history of cocaine self-administration (1.38 ± 0.16; *n* = 19 cells; *Mann-Whitney U = 45; p = 0.11^g^*; **Figure 4A**). Similarly, there were no differences in RI when the cocaine self-administering rats were grouped based on their behavioral classification as being goal-directed (1.11 ± 0.12; *n* = 11 cells) or habitual (1.76 ± 0.30; *n* = 8 cells) in relation to the yoked-saline controls (0.99 ± 0.13; *n* = 8 cells) or each other (*Kruskall-Wallis statistic = 5.02; p = 0.08^h^*; **Figure 4A**). These findings suggest that AMPA receptor subunit composition and CP-AMPA receptor insertion are likely unchanged in D2 MSNs of the DLS of rats with a history of cocaine self-administration, regardless of whether their cocaine-seeking behavior is classified as goal-directed or habitual.

**Figure 4:**
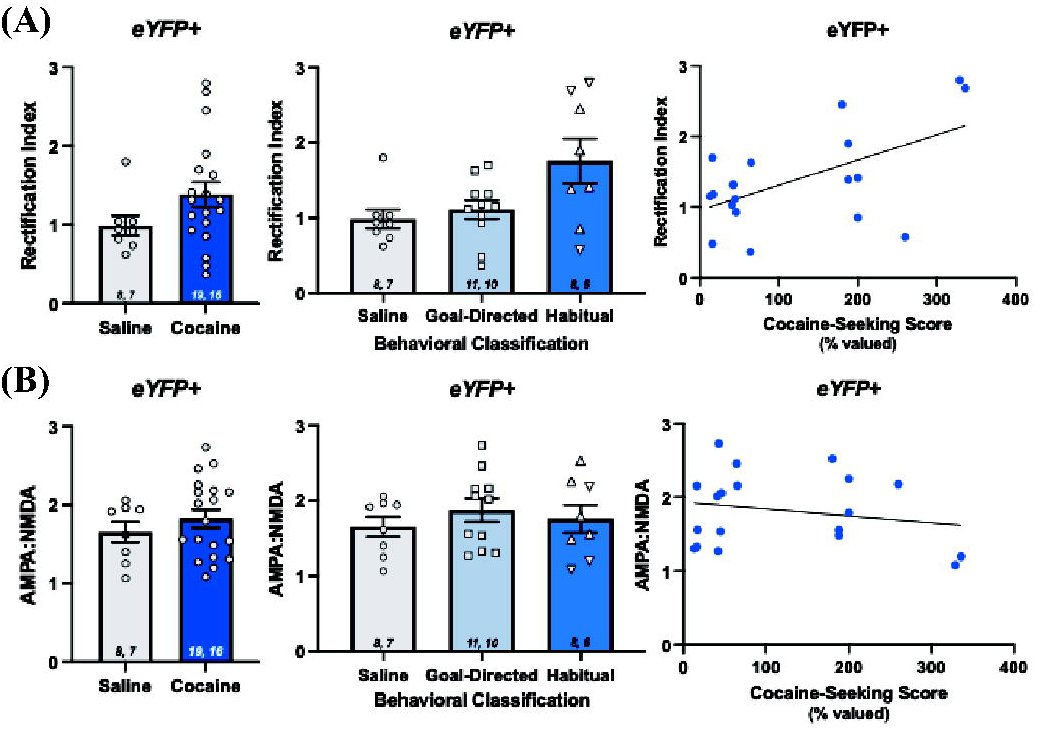
Measures of synaptic strength in eYFP+ cells of the DLS of rats with a history of cocaine self-administration. **(A)** Rectification index in brain slices from saline controls and rats with a history of cocaine self­ administration (left; *Mann-Whitney U test; p* > 0.05), and in cocaine-experienced rats characterized as goal-directed or habitual in their cocaine-seeking behavior (middle; *Kruskal-Wallis test; p* > 0.05). Correlation between rectification index and cocaine-seeking score (right; *Spearman r*= 0.35; *p* > 0.05). **(B)** AMPA:NMDA receptor ratio in eYFP+ cells of saline controls and rats with a history of cocaine self-administration (left; *unpaired t-test; p* > 0.05), and in goal-directed and habitual classifications (middle: *one-way ANOVA; p* > 0.05). Correlation between AMPA:NMDA receptor ratio and cocaine-seeking score (right; *Spearman r* = −0.004; *p* > 0.05). Data expressed as the mean ± SEM. Sample sizes for each group denoted in each corresponding bar as: number of cells, number of animals. Upside down triangles represent data points from animals trained an additional six days on Chain :#4. Correlation data fit with a linear regression line of best fit.

Increases in synaptic strength have been observed in the NAc following cocaine exposure, as well as in the DLS in the context of habitual instrumental behaviors. Therefore, we next assessed AMPA:NMDA receptor ratio as a measure of synaptic strength (Glasgow et al., 2019). AMPA receptor-mediated EPSCs were obtained at −70mV, and NMDA receptor-mediated EPSCs were obtained 50–60ms after the stimulus artifact when cells were held at +40mV (Shan et al., 2014). AMPA:NMDA receptor ratios were not significantly different between yoked-saline controls (1.65 ± 0.13; *n* = 8 cells) and rats with a history of cocaine self-administration (1.83 ± 0.11; *n* = 19 cells; *unpaired t-test; t = 0.88; p = 0.39^i^*; **Figure 4B**). Further, AMPA:NMDA receptor ratios were not significantly different between the yoked-saline controls and the cocaine-experienced rats classified as goal-directed (1.88 ± 0.15; *n* = 11 cells) or habitual (1.76 ± 0.18; *n* = 8 cells) or when compared to each other (*one-way ANOVA, F_(2, 24)_ = 0.51; p = 0.61^j^*; **Figure 4B**). Taken together, the AMPA:NMDA receptor ratio measurements in addition to the RI and EPSC analyses indicate that synaptic strength is unaltered in D2-MSNs of the DLS in rats with a history of cocaine self-administration and regardless of whether the cocaine-seeking behavior is under goal-directed or habitual control.

As we observed a wide range of cocaine-seeking scores in our animals, we also performed a correlation analysis between cocaine-seeking scores and measures of both RI and AMPA:NMDA receptor ratios **(Figure 4).** There was no significant correlation between cocaine-seeking scores and RI values in eYFP+ cells (*Spearman r = 0.3496; p = 0.14^k^*). Moreover, there was no significant correlation between cocaine-seeking scores and AMPA:NMDA receptor ratios (*Spearman r = −0.0035; p = 0.99^l^*).

### Electrophysiological recordings of MSNs not expressing eYFP reveal minimal changes in excitatory synaptic strength

Within the same brain slices from which recordings from eYFP+ cells in the DLS were obtained, we also recorded from cells *not* expressing eYFP, which could be either D1-MSNs or non-transfected D2-MSNs. Analysis of the current-voltage relationships in eYFP-cells of yoked-saline controls and rats with a history of cocaine self-administration revealed no overall changes in EPSCs (**Figure 5**). Specifically, as in the eYFP+ neurons, the current-voltage relationship between −70 and +40 mV did not display any apparent differences. (**Figure 5B**). We next assessed the RI, as described above. Again, there were no significant differences in RI between yoked-saline controls (1.72 ± 0.08; *n* = 6 cells) and rats with a history of cocaine self-administration (1.56 ± 0.09; *n* = 14 cells; *Mann-Whitney U = 31; p = 0.40^m^*; **Figure 6A**). However, comparison of the RIs across the yoked-saline controls and cocaine self-administering rats broken down into goal-directed (1.34 ± 0.1; *n* = 6 cells) and habitual (1.71 ± 0.09; *n* = 8 cells) classifications did reveal an overall significant effect of classification (*Kruskall-Wallis statistic = 6.1; p = 0.04^n^*). *Post hoc* analysis, however, failed to identify significant differences between the yoked-saline controls and either behavioral classification of cocaine-experienced rats or between cells from rats classified as either goal directed or habitual in their cocaine-seeking behavior (*Dunn’s multiple comparisons test adjusted p values; vs. goal-directed: p = 0.09, vs. habitual: p > 0.9*; goal-directed vs habitual: *p = 0.06°*; **Figure 6A**). These findings therefore suggest that there are no AMPA receptor subunit changes or insertion of CP-AMPA receptors in eYFP-cells in the DLS of rats with a history of cocaine self-administration, regardless of whether the rats are goal-directed or habitual in their cocaine-seeking behavior.

**Figure 5:**
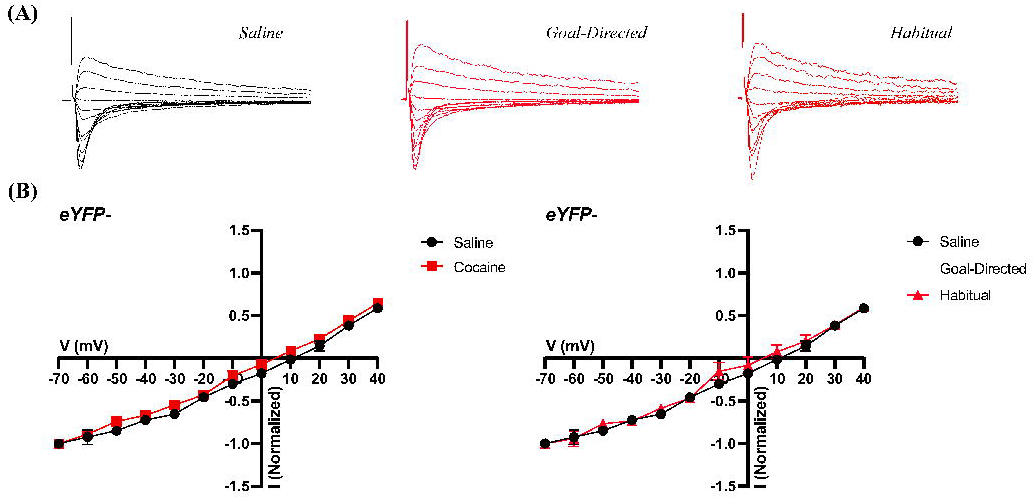
EPSCs in eYFP-cells of the DLS obtained from acute brain slices of cocaine­ experienced and yoked-saline control rats. **(A)** Representative normalized traces of evoked EPSCs in brain slices from control (left) and cocaine-experienced rats classified as goal-directed (middle) or habitual (right) in their cocaine­ seeking behavior. **(B)** IN relation of evoked EPSC responses between −70mV and +40mV in the DLS of eYFP-cells between saline *(n* = *6* cells) and cocaine-experienced rats (left; *n* = 14 cells) classified as goal-directed *(n* = *6* cells) or habitual *(n* = 8 cells) in their drug-seeking behavior (right).

**Figure 6:**
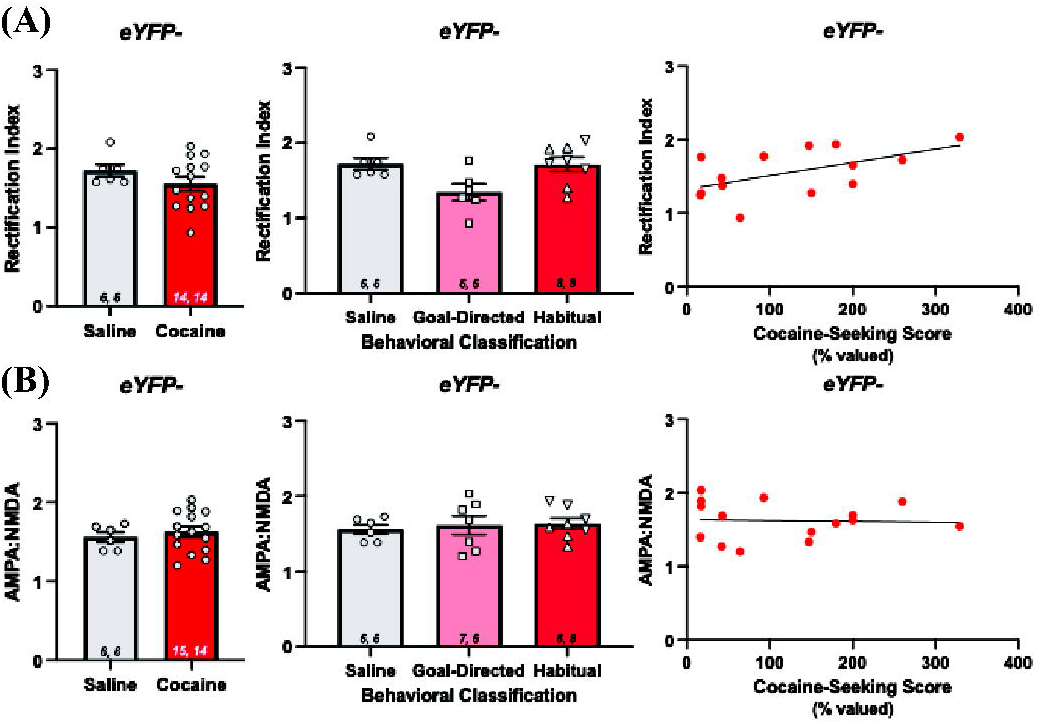
Measures of synaptic strength in eYFP-eels of the DLS of rats with a history of cocaine self-administration. **(A)** Rectification index in brain slices from saline controls and rats with a history of cocaine self­ administration (left; *Mann-Whitney U test; p* > 0.05), and in cocaine-experienced rats characterized as goal-directed or habitual in their cocaine-seeking behavior (middle; *Kruskal-Wallis test; p* = 0.05; *Dunn’s post hoc multiple comparisons adjusted p values; vs goal-directed: p* > *0.05, vs habitual: p* > *0.05, goal­ directed vs habitual: p* > 0.05). Correlation between rectification index and cocaine-seeking score (right; *Spearman r* = 0.52: *p* > 0.05). **(B)** AMPA:NMDA receptor ratio in eYFP-cells of saline controls and rats with a history of cocaine self-administration (left; *Mann-Whitney U test; p* > 0.05). and in goal-directed and habitual classifications (middle: *Kruskal-Wallis; p* > 0.05). Correlation between AMPA:NMDA receptor ratio and cocaine-seeking score (right; *Spearman r* = −0.07; *p* > 0.05). Data expressed as the mean ± SEM. Sample sizes for each group denoted in each corresponding bar as: number of cells, number of animals. Upside down triangles represent data points from animals trained an additional six days on Chain #4. Correlation data fit with a linear regression line of best fit.

Analysis of AMPA:NMDA receptor ratios in cells not expressing eYFP from saline (1.56 ± 0.06; *n* = 6 cells) and cocaine-experienced (1.63 ± 0.07; *n* = 15 cells) rats likewise revealed no significant differences in the ratios (*Mann-Whitney U = 38; p = 0.62^p^*; **Figure 6B**). Similarly, there were no differences in AMPA:NMDA receptor ratios when the rats with the history of cocaine self-administration were categorized as goal-directed (1.62 ± 0.12; *n* = 7 cells) or habitual (1.64 ± 0.07; *n* = 8 cells) in their cocaine seeking in relation to each other and the yoked-saline controls (*Kruskal-Wallis statistic = 0.31; p = 0.86^q^*; **Figure 6B**). Together, the lack of differences in the RI and AMPA:NMDA receptor ratios between yoked-saline controls and cocaine-experienced rats categorized as goal-directed or habitual in their cocaine-seeking indicate that synaptic strength in eYFP-neurons of DLS is also unaffected by a history of cocaine self-administration, regardless of whether the cocaine-seeking behavior is characterized as goal-directed or habitual.

Similar to analyses performed in eYFP+ cells, we also performed correlation analyses between cocaine-seeking scores and measures of RI values and AMPA:NMDA receptor ratios in the eYFP-cell population **(Figure 6).** We did not observe a significant correlation between cocaine-seeking scores and RI values (*Spearman r = 0.5242; p = 0.0567^r^*). Similarly, we did not observe a significant correlation between cocaine-seeking scores and AMPA:NMDA receptor ratio (*Spearman r = −0.068; p = 0.81^s^*).

Despite no significant differences observed in RI values and AMPA:NMDA receptor ratios between saline and cocaine-experienced rats, we did observe a significant difference in the RI values between eYFP+ and eYFP-neurons of the yoked saline controls (compare Figs. 4A and 6A). RI values obtained from the eYFP-cells (1.72 ± 0.08) were significantly greater than those from eYFP+ cells (0.99 ± 0.13; *Mann-Whitney U test, p* < 0.05^t^). This observation suggests that the eYFP+ and eYFP-cells are distinct cellular populations (presumed D2 vs. D1 MSNs), and eYFP-cells exhibit more pronounced inward rectification in comparison to eYFP+ cells. In contrast, AMPA:NMDA receptor ratio values did not differ between eYFP+ (1.65 ± 0.13) and eYFP-(1.56 ± 0.06^u^) cells from the saline control rats (compare Figs. 4B and 6B). Together, these findings suggest that D2 MSNs and presumed D1 MSNs display inherent differences in expression of CP-AMPA receptors but display no differences in synaptic strength under drug-naïve conditions.

### Paired pulse ratios reveal significant differences in both eYFP+ and eYFP-cells in the DLS of rats with a history of cocaine self-administration and classified as goal-directed or habitual in their cocaine-seeking behavior

Within the same recording session, we obtained PPRs as a measure of presynaptic short-term synaptic plasticity. Measures of the PPR in D2 MSNs were not significantly different between yoked-saline controls (1.41 ± 0.10; *n* = 7 cells) and rats with a history of cocaine self-administration (1.47 ± 0.1; *n* = 13 cells; *Mann-Whitney U = 37; p = 0.54^v^*; **Figure 7A**). In contrast, when PPR was assessed across rats classified as goal-directed (1.62 ± 0.09; *n* = 9 cells) or habitual (1.12 ± 0.15; *n* = 4 cells) in their cocaine-seeking behavior, the analysis revealed an overall significant effect (*Kruskal-Wallis statistic = 6.7; p = 0.03^w^*; **Figure 7A**). The *post hoc* analysis did not reveal significant differences between goal-directed or habitual rats relative to the yoked-saline controls (*Dunn’s multiple comparisons test adjusted p values; goal-directed: p = 0.37, habitual: p = 0.74^x^*; **Figure 7A**). However, a significant difference was observed in the PPRs between the goal-directed and habitual groups, with the PPR of the habitual group being significantly lower than that in the goal-directed group (*Dunn’s multiple comparisons test adjusted p value = 0.04^x^*; **Figure 7A**), suggesting paired pulse facilitation is weaker in D2 MSNs in the DLS of rats classified as habitual in their cocaine-seeking behavior than in those classified as goal-directed in their cocaine-seeking behavior. Decreased PPR is consistent with an increase in the probability of release of synaptic vesicles upon the first stimulation event from presynaptic afferent projections, reducing the number of vesicles available for release upon the next stimulation event, and thereby reducing the amplitude of the second evoked EPSC in D2 MSNs in the DLS of rats that are habitual in their cocaine-seeking behavior.

**Figure 7:**
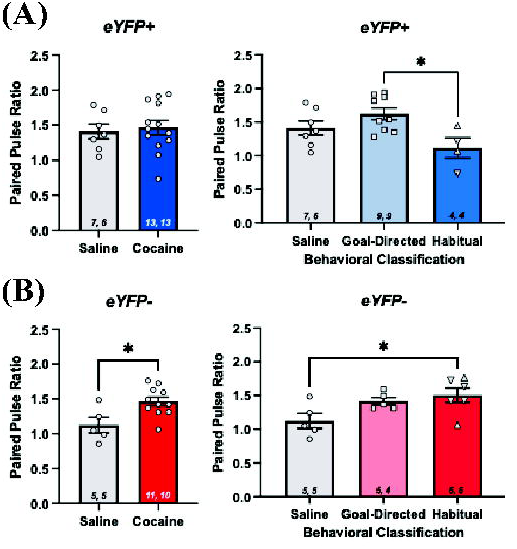
Paired pulse ratio (PPR) in the DLS of yoked-saline control and cocaine­ experienced rats classified as goal-directed or habitual in their cocaine-seeking behavior. **(A)** PPR values in brain slices from eYFP+ cells between control and cocaine-experienced rats (left; *Mann-Whitney U test; p* > 0.05), as well as between goal-directed and habitual behavioral classifications (right; *Kruskal-Wallis test with Dunn’s post hoc test, p* < 0.05). **(B)** PPR values from eYFP-cells obtained from control and cocaine-experienced rats (left; *Mann-Whitney U test; p* < 0.05), and goal-directed or habitual classifications (right; *Kruskal-Wallis test with Dunn’s post hoc test, p* < 0.05). Data expressed as mean ± SEM. Sample sizes for each group denoted in each corresponding bar as: number of cells, number of animals. Upside down triangles represent data points from animals trained an additional sex days on Chain #4.

In contrast to eYFP+ neurons, PPRs from eYFP-neurons in DLS of rats with a history of cocaine self-administration (1.46 ± 0.06; *n* = 11 cells) were significantly greater than those in yoked-saline controls (1.12 ± 0.11; n = 5 cells; *Mann-Whitney U = 7; p = 0.02^y^*; **Figure 7B**). Furthermore, analysis across yoked-saline controls and cocaine self-administering rats classified as goal-directed (1.4 ± 0.06; *n* = 5 cells) or habitual (1.5 ± 0.11; *n* = 6 cells) in their cocaine seeking revealed an overall significant effect (*Kruskal-Wallis statistic = 5.9; p = 0.04^z^;* **Figure 7B**). *Post hoc* analysis revealed that the PPRs of rats classified as habitual, but not goal-directed, in their cocaine seeking behavior were significantly greater than those in eYFP-neurons from the yoked-saline controls (*Dunn’s multiple comparisons test adjusted p values; goal-directed: p = 0.33, habitual: p < 0.05^aa^*; **Figure 7B**). Together, the PPR values recorded from eYFP-neurons in the DLS suggest the possibility that a history of cocaine seeking and taking leading to habitual control over drug-seeking behavior is associated with a decrease in the initial probability of presynaptic glutamate release onto the eYFP-neurons.

## Discussion

The transition from goal-directed to habitual control over drug-related behavior is thought to play a critical role in the transition between recreational drug use and addiction. While much is known about cell-type-specific changes in synaptic strength in the DLS in the development of habitual behavior, specifically increasing synaptic strength of D2 MSNs (Yin et al., 2009; O’Hare et al., 2016; Lipton et al., 2019), relatively little is known about the state of excitatory synaptic function in MSNs of the DLS in the context of habitual cocaine-seeking behavior. Therefore, the present study was designed to investigate changes in synaptic strength induced by a history of cocaine self-administration and in relation to goal-directed vs habitual control over cocaine-seeking behavior. Overall, we found that synaptic strength, either through changes in the insertion of CP-AMPA receptors as reflected in changes in rectification index or in AMPA:NMDA receptor ratios, is unaffected in eYFP+ D2 MSNs and in other unlabeled MSNs by a history of cocaine self-administration under a seeking-taking paradigm and regardless of whether the cocaine-seeking is characterized as goal-directed or habitual. We did, however, observe evidence of short term plasticity, as measured by PPRs, with there being a decrease in PPRs in eYFP+ neurons from rats whose cocaine-seeking was classified as habitual as compared to those classified as goal-directed and an increase in PPRs in eYFP-neurons from cocaine-experienced rats, specifically those classified as habitual in their cocaine-seeking behavior. Together, the present study represents the first assessment of cell-type-specific adaptations in excitatory synaptic transmission in MSNs of DLS in the context of habitual control over cocaine-seeking behavior and provides a foundation for future inquiries into the synaptic perturbations induced by repeated cocaine use.

The present study capitalized on a cocaine self-administration paradigm previously established and replicated in the literature by Zapata et al. (2010) and Giangrasso et al. (2023). This paradigm reliably produces rats whose cocaine-seeking behavior can be classified as goal-directed (cocaine-seeking score of ≤70%), intermediate (cocaine-seeking score between 71% – 79%) or habitual (cocaine-seeking score of ≥80%) based on sensitivity to devaluation of the cocaine-taking lever via outcome omission. Zapata et al. (2010) reported rats with goal-directed cocaine-seeking behavior transitioned to habitual control with extended training. Similarly, to increase the number of rats meeting criterion for being classified as habitual in their cocaine-seeking, a subset of our rats were run an additional six days of Chain #4 prior to devaluation training. Of this subset, 60% were classified as habitual, whereas only 27% of those trained on the original six days of Chain #4 were classified as habitual. These findings complement those of Zapata and colleagues (2010) who also showed that extended training on the chained self-administration paradigm led to a greater percentage of rats exhibiting habitual control over their cocaine seeking. The present results also extend those findings by demonstrating that simply extending the time on the Chain #4 schedule, without taking the animals through extinction and testing first, is a viable approach to promote development of habitual control over cocaine-seeking behavior in this paradigm.

A viral vector under a D2-specific promotor was used to express eYFP for patch clamp studies, as this has been shown to reliably infect D2 MSNs in ventral striatum (Zalocusky et al., 2016; Soares-Cunha et al., 2022). Importantly, RNAScope™/IHC analysis indicated that eYFP expression in our hands using this viral construct was limited to Penk1-expressing MSNs and was not observed to colocalize in Tac1-expressing MSNs, supporting the claim that eYFP+ labeled neurons are D2 MSNs (Figure 2). As cholinergic interneurons have also been shown to express D2 receptors, it is possible some cells labeled were not MSNs (Kreitzer & Malenka, 2008). However, in all electrophysiological recordings, intrinsic properties of the recorded cells, as well as visual cues, indicated these were MSNs rather than cholinergic interneurons. Specifically, cholinergic interneurons display depolarized resting membrane potentials (−54mV to −60mV) and are clearly identifiable within the slice by their large cell body in comparison to MSNs (Kawaguchi, 1992; Bennett & Wilson, 1999). Further, we did not observe any eYFP+ cells that did not also express Penk1 in our RNAScope™/IHC analysis, suggesting eYFP expression is restricted to MSNs, as cholinergic interneurons in the striatum do not express Penk1 (Meredith & Chang, 1994). Together, electrophysiological recordings and RNAScope™/IHC analysis suggest the eYFP+ cells in the DLS are D2 MSNs. The specific phenotype of the eYFP-neurons from which we recorded is unknown. However, these cells displayed characteristics consistent with MSNs, indicating that these neurons are likely either D1 MSNs or non-transfected D2 MSNs. Our observations that the RI and PPRs differ between the eYFP+ and eYFP-neurons further suggests that these eYFP-cells are likely a separate population of MSNs (i.e., D1 MSNs).

The present findings suggest excitatory synaptic parameters of the eYFP+ D2 MSNs are unaffected by cocaine self-administration experience whether cocaine-seeking behavior is under goal-directed or habitual control. In the analysis of RI and AMPA:NMDA receptor ratio in eYFP+ cells, there were no significant differences observed between yoked-saline control rats and rats with a history of cocaine self-administration, whether goal-directed or habitual in their cocaine seeking. This lack of any differences in the RI and AMPA:NMDA ratio indicate that the number and composition of postsynaptic AMPA receptors in D2 MSNs are not changed by cocaine experience (Figure 4). Similarly, in eYFP-cells, there were no significant differences in RI or AMPA:NMDA receptor ratio, indicating postsynaptic AMPA receptors are also unaltered in this cell population, presumed to be D1 MSNs (Figure 6).

The lack of changes in excitatory post-synaptic measures in DLS of rats with a history of cocaine self-administration was surprising given that rats clearly differed in the nature of the behavioral control over their cocaine seeking (Figure 1). Numerous studies of dorsal striatum have demonstrated synaptic changes associated with the acquisition and expression of goal-directed and habitual behaviors (e.g., Amaya & Smith, 2018; Lipton et al., 2019). For example, increases in AMPA:NMDA receptor ratio in D1 MSNs of the DMS is observed in goal-directed behaviors (Shan et al., 2014). However, in habitual behaviors, D2 MSNs of the DLS exhibit similar robust synaptic changes (Yin et al., 2009; O’Hare et al., 2016). As such, we expected to observe similar synaptic changes in DLS of our habitual cocaine-seeking rats, yet we did not. Cocaine is known to influence cell-type-specific synapses in a projection-specific manner (Zinsmaier et al., 2022). For example, within the NAc, cocaine self-administration differentially affects synapses from the infralimbic prefrontal cortex (ILC-NAc) and the ventral hippocampus (vHipp-NAc) onto D1 MSNs, with ILC-NAc synapses enriched with CP-AMPA receptors and vHipp-NAc synapses enriched with calcium-impermeable AMPA receptors (Pascoli et al., 2014; Zinsmaier et al., 2022). Therefore, it is conceivable that the lack of differences observed in the present study reflects a similar mechanism of cocaine-induced, afferent-specific changes in the dorsal striatum. Such differences would be obscured in our studies because the local stimulation used did not activate specific afferent pathways. Additional experiments will be needed to identify any pathway-specific changes in excitatory synaptic transmission onto MSNs of the DLS using afferent-selective stimulation approaches, such as with optogenetics. Moreover, as mentioned above, identified D1 MSNs remain to be more thoroughly interrogated, as D1 MSNs in DLS also exhibit activity changes as behavior is learned and expressed, suggesting similar synaptic changes are occurring to D1 MSNs (Smith et al., 2021). It is therefore conceivable that D1 MSN synapses in the DLS may exhibit synaptic strengthening in cocaine-experienced animals and habitual cocaine-seeking behavior.

Despite not seeing differences in synaptic strength in eYFP+ D2 MSNs and eYFP-negative MSNs in DLS, we did observe statistically significant differences in short-term plasticity, as measured by PPRs. Specifically, there was weaker paired-pulse facilitation of input to D2 MSNs from rats classified as habitual in their cocaine-seeking relative to rats classified as goal-directed. Further, in the eYFP-negative MSNs, paired-pulse facilitation was stronger in the rats with a history of cocaine self-administration, namely in the eYFP-negative MSNs from rats characterized as habitual in their cocaine-seeking relative to yoked-saline controls. Paired pulse facilitation has long been considered a form of short-term synaptic plasticity related to the probability of neurotransmitter vesicle release at the synapse, and is, thereby, largely considered to be presynaptic in origin (Cepeda et al., 2008; Glasgow et al., 2019). This classical view of the PPR would suggest, therefore, differential changes in release probability onto D2 MSNs and presumed D1 MSNs in the DLS in rats with habitual control over drug-seeking behavior. However, there are numerous other mechanisms that may contribute to alterations in PPR (Glasgow et al., 2019). For example, some studies have shown that PPR values stay constant as the release probability is altered (Manita et al., 2007; Deutschmann et al., 2021), and seminal work on striatal MSNs has shown dissociation between PPR and other measures of release probability onto D2 MSNs (Cepeda et al., 2008). It is also important to acknowledge that sample sizes from our PPR analyses are small (4-9 cells/group) and the resulting power (0.77 for eYFP+ analyses, 0.73 for eYFP-analyses) slightly below the typical desired power of 0.8. Therefore, the present PPR data suggest the possibility that short-term plasticity in excitatory afferents onto MSNs in DLS may be altered in rats that have developed habitual control over their drug-seeking behavior, and the nature of those alterations may differ between D1 and D2 MSNs. That said, the present data should be considered preliminary and future work should more thoroughly examine whether differences in short-term plasticity of excitatory input onto D1 and D2 MSNs in DLS are apparent in rats with habitual control over cocaine-seeking behavior.

It is important to note a limitation to the study of RI values in our preparation. Specifically, the preparations in which the RI in eYFP+ and eYFP-cells were determined did not include NMDA-receptor antagonists, because we were attempting to maximize data collection from each rat, given the duration of the training required. Consequently, there is the possibility that our RI measures may be confounded by NMDA receptor-mediated currents at depolarized potentials (+40mV). While AMPA and NMDA receptors display different kinetic properties, with NMDA receptors possessing notably slower rates of synaptic transmission, NMDA receptor-mediated currents at depolarized potentials may still contribute to synaptic responses and, thereby, contaminate the outward AMPA EPSC measured at +40 mV. Importantly, many aspects of NMDA receptors can contribute to rectification patterns, including NMDA receptor subunit composition and extrasynaptic NMDA receptor activation (Sun & Liu, 2007; Jeun et al., 2009). It is possible that NMDA receptors containing the NR2A subunit, which possess faster kinetics than those containing the NR2B subunit, may contribute to the outward EPSC at depolarized potentials (Jeun et al., 2009). To this point, NR2A subunits are expressed at higher levels in DLS of adult rodents and NMDA receptor-mediated currents in the DLS exhibit faster rise times and decay constants relative to ventromedial striatum (Standaert et al., 1994; Chapman et al., 2003). Given this limitation, future studies should assess RI in the presence of APV, an NMDA receptor antagonist, or through the use of the specific CP-AMPA receptor antagonist, Naspm.

This limitation regarding RI notwithstanding, differences in the RI of eYFP+ vs. eYFP-neurons were apparent in the yoked saline controls, with the RI being ∼70% greater in the eYFP-MSNs. Importantly, the RI in the eYFP+ MSNs of the saline controls was ∼1, suggesting no inward rectification and, thus, few CP-AMPA receptors, consistent with expression of the GluR2 subunit in all identified striatal efferent neurons (Deng et al., 2007). The present observation, however, suggests the possibility that eYFP-MSNs—presumed D1 MSNs—in DLS may express more CP-AMPA receptors relative to D2 MSNs (eYFP+ MSNs). While differential changes in CP-AMPA expression in D1 vs. D2 MSNs have been reported in the nucleus accumbens in the context of incubation of cocaine craving (c.f., Wolf 2016; Terrier et al., 2016), to our knowledge no one has heretofore reported differences in RIs in D1 vs D2 MSNs in nucleus accumbens or dorsal striatum. The present findings therefore raise the intriguing possibility that the eYFP-MSNs (presumed D1 MSNs) in DLS express more CP-AMPA receptors. Of course, the caveat regarding the potential role of NMDA receptor subunit composition still pertains, although to date differences in the expression of NMDA receptor subunits (Standaert et al., 1999; Ganguly & Keefe, 2001) or excitatory synaptic NMDA receptor subtypes between D1 and D2 MSNs have also not been reported (Krajeski et al., 2019). Clearly, additional studies are needed to thoroughly examine AMPA receptor subtype expression and function between these two populations of MSNs in the DLS.

The present study represents the first examination of whether there are MSN-specific adaptations in excitatory synaptic transmission in the DLS of rats classified as goal-directed or habitual in their cocaine-seeking behavior. Overall, we observed that excitatory synaptic strength in D2 MSNs as well as unlabeled, presumed D1 MSNs is not different in rats characterized as habitual vs. goal-directed in their cocaine-seeking behavior, either relative to each other or to yoked-saline controls. Therefore, there do not appear to be changes in synaptic strength assessed through AMPA:NMDA receptor ratio or RI in MSNs of the DLS associated with the development of habitual control over drug-seeking behavior. Interesting preliminary data, however, suggest the possibility that changes in short-term plasticity, as evidenced by changes in PPRs, may be associated with the development of habitual control over behavior and that such changes may differ between D2 and D1 MSNs. While cell-type-specific adaptations have been well-characterized in the NAc, studies of the cell-type-specific synaptic changes of the dorsal striatum in drug-related behaviors remains in its infancy. Nevertheless, the present study represents a critical starting point for future investigations to the neurobiology of the dorsal striatum in the development and progression of cocaine use behaviors.

## Notes

### Competing Interest Statement

The authors have declared no competing interest.

### Funding

- NIH: R21DA0466000
- University of Utah: T32NS115664
- Skaggs Graduate Research Fellowship

